# Comparison of ultracentrifugation and a commercial kit for isolation of exosomes derived from glioblastoma and breast cancer cells

**DOI:** 10.1101/274910

**Authors:** Frøydis Sved Skottvoll, Henriette Engen Berg, Kamilla Bjørseth, Kaja Lund, Norbert Roos, Sara Bekhradnia, Bernd Thiede, Cecilie Sandberg, Einar Osland Vik-Mo, Hanne Roberg-Larsen, Bo Nyström, Elsa Lundanes, Steven Ray Wilson

**Affiliations:** Department of Chemistry, University of Oslo, Post Box 1033, Blindern, NO-0315 Oslo, Norway; Department of Microbiology, Unit Cell Signaling, Oslo University Hospital, Gaustadalleen 34, NO-0372 Oslo, Norway; Department of Biosciences, University of Oslo, Post Box 1066, Blindern, NO-0316 Oslo, Norway; Vilhelm Magnus Laboratory of Neurosurgical Research, Institute for Surgical Research and Department of Neurosurgery, Oslo University Hospital, 4950 Nydalen, NO-0424 Oslo, Norway; Institute of Clinical Medicine, Faculty of Medicine, University of Oslo, Post Box 1171, Blindern, 0318 Oslo, Norway

**Keywords:** Exosomes, Ultracentrifugation, Proteomics, Glioblastoma, Breast cancer, LC-MS/MS

## Abstract

Exosomes are a potentially rich source of biomarkers, but their isolation and characterization can be challenging. For isolation of exosomes, differential ultracentrifugation (a traditional approach) and an isolation kit from a major vendor (Total Exosome Isolation Reagent from Thermo Fisher Scientific) were compared. “Case study” exosomes were isolated from cell culture media of two different cell sources, namely patient-derived cells from glioblastoma multiforme and the breast cancer cell line MDA-MB-231. For both isolation methods, transmission electron microscopy and dynamic light scattering indicated the presence of exosomes. The kit- and UC isolates contained similar amounts of protein measured by the bicinchoninic acid (BCA) assay with absorbance at 562 nm. Using western blot, positive exosome markers were identified in all isolates. Potential biomarkers for both diseases were also identified in the isolates using LC-MS/MS. However, WB and LC-MS/MS also revealed negative exosome markers regarding both isolation approaches. The two isolation methods had an overall similar performance, but we hesitate to use the term “exosome isolation” as impurities may be present with both isolation methods. LC-MS/MS can detect disease biomarkers in exosomes and is also highly useful for critical assessment of exosome enrichments.

## 1 Introduction

Exosomes are extracellular vesicles (EVs) with membrane-surrounded bodies which are secreted from cells to the extracellular environment as a part of the endocytic pathway [1]. Exosomes are formed by invagination of an endosome membrane to create intraluminal vesicles inside the endosome, i.e. multivesicular bodies (MVBs), and are secreted when the endosomes fuse with the plasma membrane [2]. Exosomes commonly contain proteins originating from the cellular cytosol and the plasma membrane, nucleic acids (e.g. DNA, mRNA, microRNA and non-coding RNA), lipids and metabolites [3–5,1,6–8], and are believed to take part in e.g. cell-cell communication, transfer of proteins/nucleic acids, coagulation and antigen presentation [6,9].

Cancer cells have been found to release more exosomes than stromal cells [10,11] and exosomes are associated with metastasis and tumor progression [7,12,13]. Hence, cancer exosomes may be a source of biomarkers for diagnosing cancers such as breast cancer (BC) and glioblastoma multiforme (GBM) when e.g. isolated from body fluids. BC is the predominant type of female cancer [14], with recurrent metastatic disease being responsible for the majority of BC-caused deaths [15]. GBM is the most frequent and malignant form of brain cancer [16–18]. The diagnosis of both BC and GBM rely on highly invasive patient tissue biopsies at relatively late stages [16,19,20]. Thus, a non-invasive disease monitoring is desirable for both BC and GBM, and can be achieved by measuring biomarkers in accessible body fluids, such as blood (liquid biopsy), for early diagnosis and prognosis assessment [16,21–23]. Hence, the isolation of exosomes for cancer biomarker discovery has emerged as an alternative to invasive methodologies [24–31,23].

Isolation of exosomes is predominantly performed from body fluids (e.g. blood, urine, and saliva) or cell culture media by centrifugation-based methods, e.g. sucrose density gradient centrifugation or ultracentrifugation (UC) [32,33]. However, common drawbacks of using UC-based exosome isolation methods are the large amounts of starting material needed, low yield, and poor reproducibility [34,35]. Moreover, there is a great need for exosome isolation protocols tailored towards smaller starting volumes for e.g. miniaturized cell culture models like organoids and “organ on a chip” [36,37]. Other exosome isolation protocols and principles have been developed to overcome the drawbacks of UC based methods. Among these, filtration, immunoaffinity capturing, size exclusion chromatography, flow field-flow fractionation and also acoustic trapping have been attempted [34,38-42,8,43,44]. In addition, different commercial exosome isolation kits are available (e.g. ExoQuick™ from Systems Biosciences, and Total Exosome Isolation^™^ from Thermo Fisher), enabling simple isolation of exosomes from small starting volumes from a wide range of matrices. The exosome isolation kits are known to be based on exosome precipitation at low-speed centrifugation after sample incubation with water-excluding polymers such as polyethylene glycol (PEG) [45].

We have compared two exosome isolation methods, namely UC and a commercial kit for precipitation of exosomes. The methods were evaluated using the following characterization techniques: WB, transmission electron microscopy (TEM), dynamic light scattering (DLS), quantitative total protein analysis using UV-Vis spectrophotometry and LC-MS/MS. “Case study” exosomes were isolated from cell culture media from free-floating patient-derived primary cell cultures from GBM biopsies (T1018) and a serum cultivated, adherently growing BC cell line (MDA-MB-231).

## 2 Materials and Methods

### 2.1 MDA MB-231 cell culturing

The BC cell line was purchased from American Type Culture Collection (ATCC, Sesto San Giovanni, Milan, Italy) and is derived from a triple-negative human metastatic breast carcinoma. The cells were maintained in Rosewell Park Memorial Institute (RPMI) 1640 growth medium depleted of phenol red (Sigma-Aldrich, St. Louis, MO, USA) supplemented with 10 % exosome-depleted fetal bovine serum (FBS) (System Biosciences, Palo Alto, CA, USA) and 1 % penicillin/streptomycin (Sigma-Aldrich). The cells were incubated in a humidifying atmosphere at 5 % CO2 and at 37 C. Prior to exosome isolation, 1-2.3 million cells (in T75-T175 culturing flasks) were incubated for 6-7 days (always using a passage lower than 12). The incubated cell culture medium was centrifuged at 906 × *g* (30 minutes at 23 °C). See also **Supplementary 1 (S1)**.

### 2.2 Glioblastoma cell culturing

The GBM cells (T1018) were derived from biopsies from a primary GBM tumor, obtained after informed consent through a biobank approved by the Regional Ethical Authorities operated at Oslo University Hospital (2016/1791). The cells were maintained in Dulbecco’s modified eagle medium with nutrient mixture F-12 (DMEM/F12, Gibco, Thermo Fisher Scientific, Waltham, MA, USA), supplemented with HEPES buffer (10 mM) and penicillin/streptomycin (100 U/mL) from Lonza (Basel, Switzerland), B27 without vitamin A (1/50) from Thermo Fisher Scientific, epidermal growth factor (20 ng/mL) and basic fibroblast growth factor (10 ng/ mL) from R&D Systems (Minneapolis, MN, USA) and heparin (2.5 μg/mL) obtained from LEO Pharma AS (Ballerup, Denmark). Under these culturing conditions, cells express stem cell markers *in vitro,* differentiate upon removal of growth factors and give rise to diffusely infiltrative tumors upon xenografting [46]. The cells were incubated in a humidifying atmosphere at 5 % CO_2_ and 37 °C in T25 flasks (Thermo Fisher Scientific). Prior to exosome isolation, the incubated cell culture medium was centrifuged twice at 453 × *g* and 1811 × *g* for 5 minutes each. The cell pellets were harvested for WB analysis. See also **S1**.

### 2.3 Exosome isolation by ultracentrifugation

For the BC and GBM cells, 9-12 mL and 60 mL cell culture media were used for centrifugation, respectively. Cell culture media were first centrifuged at 1811 × *g* (5 minutes at 20 °C). The supernatants were then centrifuged at 20 000 × *g* (20 minutes at 4 °C) with an Allegra 25R centrifuge (with TA-14-50 rotor) from Beckman Coulter (Brea, CA, USA) and the supernatants were transferred to polycarbonate ultracentrifugation tubes (Beckman Coulter) and diluted with PBS (~60 mL in each). The tubes were centrifuged twice at 100 000 × *g* (90 minutes at 4 °C) with an L-80 ultracentrifuge (45 Ti rotor) from Beckman Coulter. The supernatants were removed (leaving suspension 1 cm above the pellets) and the pellets were suspended with PBS between the centrifugations. Upon centrifugation, the supernatants were discarded and the exosome pellets (UC isolates) were suspended in either PBS (3 mL for DLS- and 50-100 μL for TEM analysis) or the preferred lysis buffer.

### 2.4 Exosome isolation by isolation kit

The isolation of exosomes with the kit was performed with the Total Exosome Isolation Reagent (from cell culture media) from Thermo Fisher Scientific (catalog no. 4478359). The isolation was performed according to the protocol of the supplier [47]. Starting volumes ranged from 0.5 mL to 9 mL cell culture medium for the BC cells and 5 mL to 6 mL for the GBM cells. The samples were centrifuged with the Allegra 25R centrifuge, and the exosome pellets (kit isolates) were suspended as with UC.

### 2.5 Protein extraction

Cell and exosome protein extracts were made by lysis with RIPA- or Nonidet™ P40 (NP40) buffer (both from Thermo Fisher Scientific) containing protease inhibitors (Protease Inhibitor Cocktail Tablets, Roche, Basel, Switzerland) and phosphatase inhibitors (PhosStop Tablets, Sigma-Aldrich). See also **S2**.

### 2.6 UV-Vis spectrophotometry

The protein amount was measured using Pierce™ BCA protein Assay Kit (Thermo Fisher Scientific), by measuring the absorbance at 562 nm. See also **S3**.

### 2.7 Western blotting

For information about WB antibodies, procedures and equipment, see **S4.**

### 2.8 Immunogold labeling and transmission electron microscopy

Samples were visualized with a JEM-1400Plus transmission electron microscope from JEOL (Tokyo, Japan) and images were recorded at 80 kV. See also **S5.**

### 2.9 Dynamic light scattering

The DLS experiments were conducted with the aid of an ALV/CGS-8F multi-detector version compact goniometer system, with 8 fiber-optical detection units, from ALV-GmbH, Langen, Germany. See S6 for more details.

### 2.10 LC-MS/MS analysis

LC-MS/MS was performed using Q-Exactive mass spectrometers (Thermo) coupled with liquid nano chromatography. Samples were prepared by in-solution and in-gel protease digestion. See **S7-9** for additional information related to LC-MS/MS analysis.

## 3 Results and Discussion

### 3.1 Similar content of protein measured in kit- and UC isolates

The protein amount per million cells (hereafter referred to as protein amount) in the BC-(**Figure 1A**) and GBM-(**Figure 1B**) isolates was measured using UV-Vis spectrophotometry. The measurements for kit isolates were 15-28 times higher than for UC isolates. A higher protein amount in exosomes isolated by the kit compared to that by UC was also observed in a study by Van Deun et al. who compared UC to the same isolation kit used in the present study for MCF7 derived exosomes [48]. However, we observed that the measured absorbance in the kit blanks was high in comparison to UC blanks, where the absorbance was below the limit of quantification. The high absorbance from the kit blanks was further assessed to establish possible UV-absorbents or scattering components in the kit reagent. However, no absorbance was measured in the kit reagent using the same protocol (i.e. absorbance at 562 nm after BCA-reaction) as for the isolates and blanks, and NMR spectroscopy showed sharp peaks implying an absence of relaxation-perturbing components, e.g. particles (results not shown). The high absorbance in the kit blanks might therefore indicate co-precipitation of proteins or other UV-absorbing compounds from the blank media. When correcting for the blank (subtracting the protein amount measured in blank samples from the protein amount in exosome isolates), the measured protein content for exosomes isolated by the kit and UC was similar.

**Figure 1.**
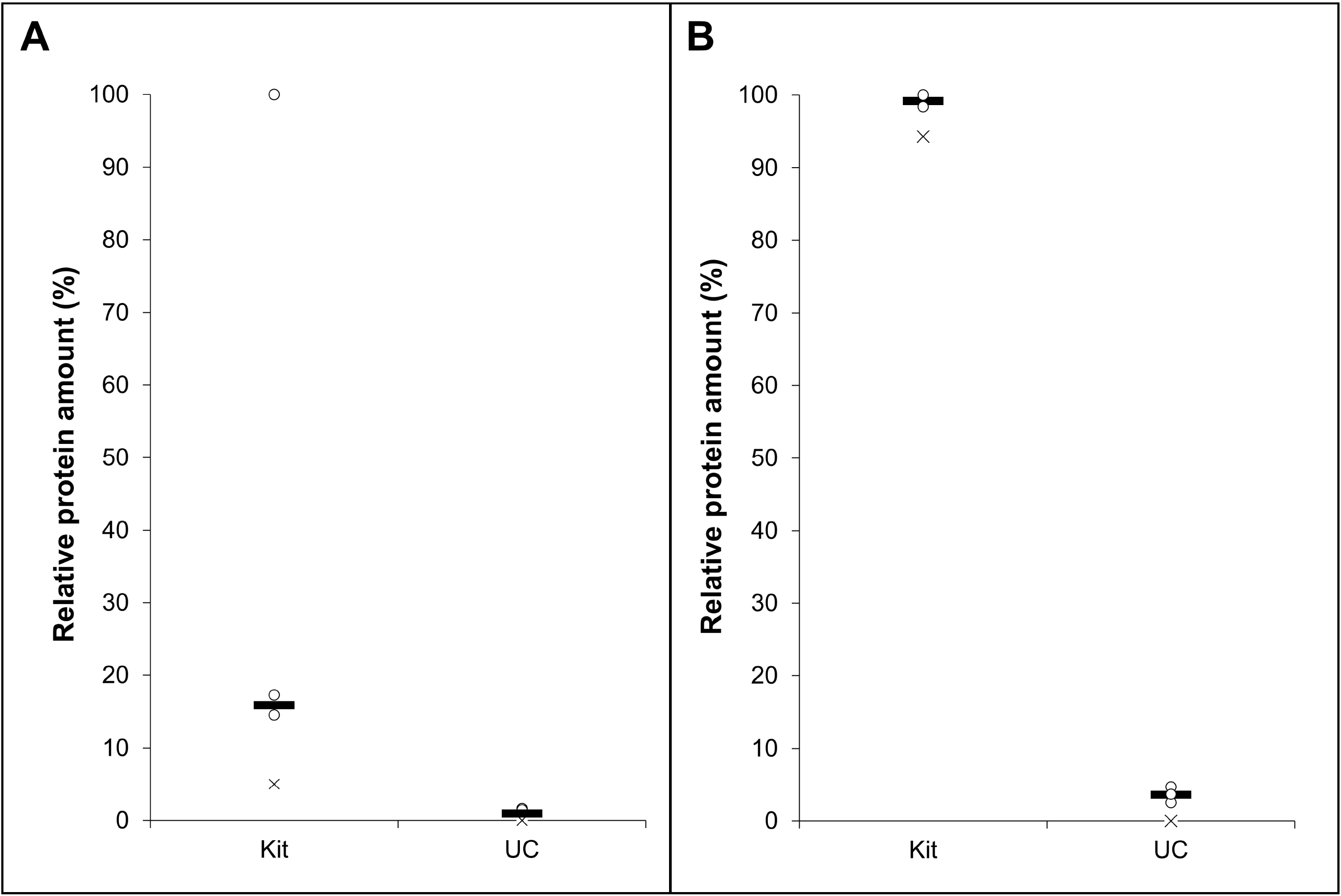
Measured relative protein amount pr. million cells in exosome samples from GBM- and BC cells isolated by kit and UC (n ≥ 2). **A**) The measured relative protein amount (%) for the BC exosome isolates. **B**) The measured relative protein amount (%) for the GBM exosome isolates. Each replicate is depicted as circles, and the median depicted as a line. The X-mark shows the measured relative protein amount in the blank sample (isolated cell culture medium). The protein amounts were measured by UV-Vis spectrophotometry (absorption at λ= 562 nm) after reaction with BCA kit reagents.

### 3.2 TEM and DLS detected vesicles in the expected size range for exosomes

Morphological analysis of the exosome samples was performed using TEM. In addition, the hydrodynamic particle size distribution was measured using DLS. Clusters of vesicles were observed in the micrographs of the samples isolated with both kit and UC (**Figure 2, AI and AIII**). Vesicle structures similar to that described in literature were observed [49,50,6]. Regarding GBM exosomes: With TEM, the UC isolates presented somewhat more distinct double membranes compared to the kit isolates. The blank samples for both isolation methods did not display membrane structures (**Figure 2, AII** and **AIV**). The DLS-analysis of the GBM isolates exhibited particles of similar sizes of 51 and 73 nm (mean) with both isolation methods (**Figure 2B**). Thus, both isolation methods gave rise to comparable exosome populations. Regarding BC exosomes: Clusters of vesicles were also in here observed in the micrographs of the samples isolated with both kit and UC (**Figure 2, CI and CIII**). Blank isolates displayed contaminations (**Figure 2, CII and CIV**), e.g. exosome-resembling vesicles were found in the UC blank using TEM (red dashed circles), and the kit blank displayed 67 nm (mean) contaminations when using DLS (**Figure 2D**). The DLS analysis also presented two distinct particle diameters in kit isolates (28 and 95 nm, mean values) while only one particle diameter was present in UC isolates (137 nm, mean value), indicating some differences in the mean particle sizes isolated with the two isolation methods. However, the sizes observed with DLS correlates well with that found in other studies (30–250 nm) [51,52,13,53,54,48,55]. Overall, the isolates showed structures resembling those of EVs, but some blanks were not entirely devoid of vesicles or particles.

**Figure 2.**
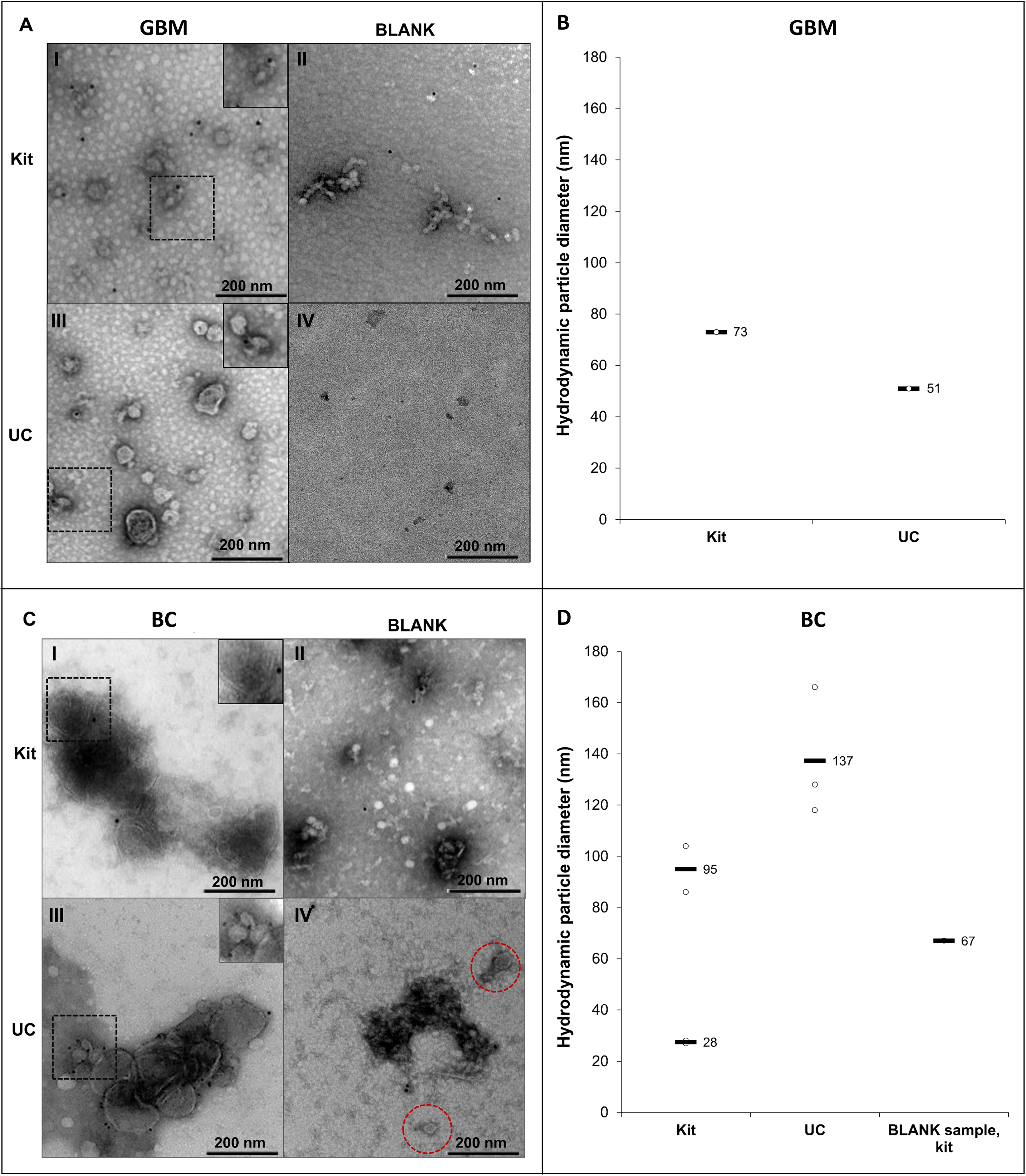
Transmission electron micrographs and hydrodynamic particle size (nm) distribution by DLS analysis of exosomes isolated by kit and UC from GBM- and BC cells. Images were taken with a magnification of 400 000, and the dashed areas were additionally zoomed. **A**) Micrographs of GBM exosome isolates (not CD9-labelled). **I** depict the micrograph from a kit isolate, **II** the kit blank, **III** a UC isolate, and **IV** the UC blank. **B**) DLS analysis of GBM exosomes isolated by kit and UC (n = 1). No particles were detected in the UC blank (n = 1). DLS analysis of the kit blank was not performed. **C**) Micrographs of BC exosome isolates (successfully CD9-labelled). **I** depict the micrograph from a kit isolate, **II** the kit blank, **III** a UC isolate, and **IV** the UC blank. **D**) DLS analysis of BC exosomes isolated by kit (n = 2) and UC (n = 3), including the kit blank (n = 1). No particles were detected in the UC blank.

### 3.3. Western blot analyses indicated the presence of exosomes for all samples

WB was performed using antibodies for a selection of positive exosome markers, namely the tetraspanins CD81, CD9 and CD63, TSG101 and flotillin-1. Calnexin was selected as a negative marker for purity evaluation as recommended by the International Society of Extracellular vesicles (ISEV) [56]. This protein is located at the endoplasmic reticulum (ER) and is assumed to signalize ER-contamination. For the GBM cells and exosomes, positive and negative exosome markers were detected in isolates from both the kit and UC (**Figure 3**). For the BC cells and exosomes, positive markers TSG101, flotillin-1 and CD9 (barely visible in the UC isolates) were detected using both isolation methods, and calnexin was not detected. The positive markers overall demonstrate the presence of exosomes in the isolates obtained using both methods, but the GBM samples could contain impurities.

**Figure 3.**
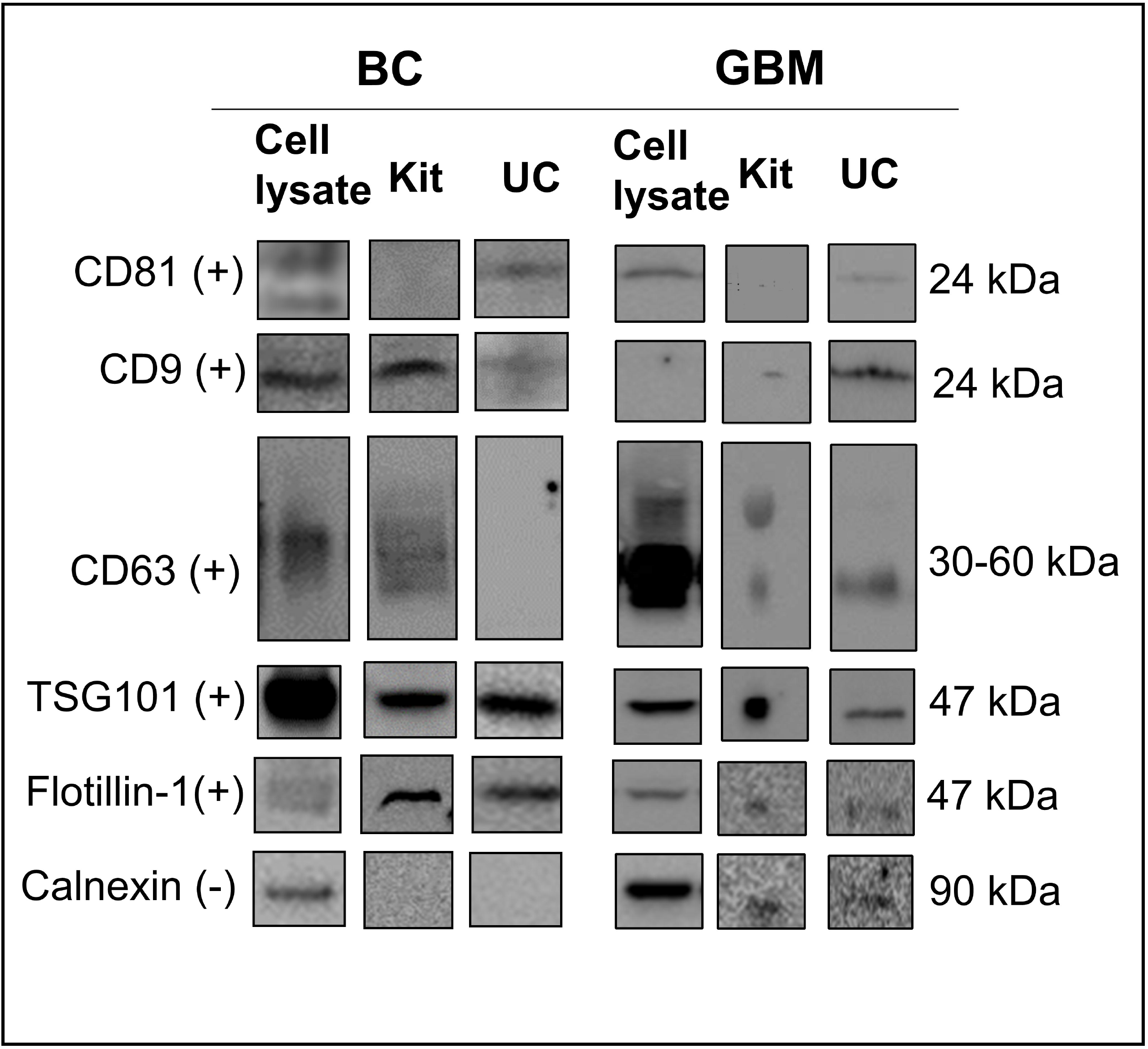
Western blot of common protein exosome markers. The protein markers CD81, CD9, CD63, TSG101, flotillin-1 (positive markers, +) and calnexin (negative marker, -) were targeted in cell lysates and exosomes isolated by kit and UC (n ≥ 2). Monoclonal mouse antibodies were used for CD81, CD9, CD63, flotillin-1 and calnexin, while a polyclonal rabbit antibody was used for TSG101. For the BC exosomes, 15 μg protein was loaded for kit isolates and 3 μg for UC isolates. For the GBM exosomes, ~14 μg was loaded for kit isolates and ~8 μg for UC isolates. Uncropped western blots are presented in **Supplemental Western Blots**.

### 3.4 LC-MS/MS studies reveal impurities, and biomarkers

The absence of calnexin (see above) in BC exosomes from both isolation methods indicates that the isolates are not contaminated with the ER. However, general proteins related to e.g. the nucleus, Golgi apparatus, mitochondrion, and ER were identified in the BC exosomes using LC-MS/MS and gene ontology (GO) annotations (**Figure 4**). Hence, untargeted LC-MS/MS suggested the presence of impurities also in the BC samples. Proteins related to the nucleosome, Golgi apparatus, mitochondrion, and ER were also identified by GO-annotation in the GBM isolates.

**Figure 4.**
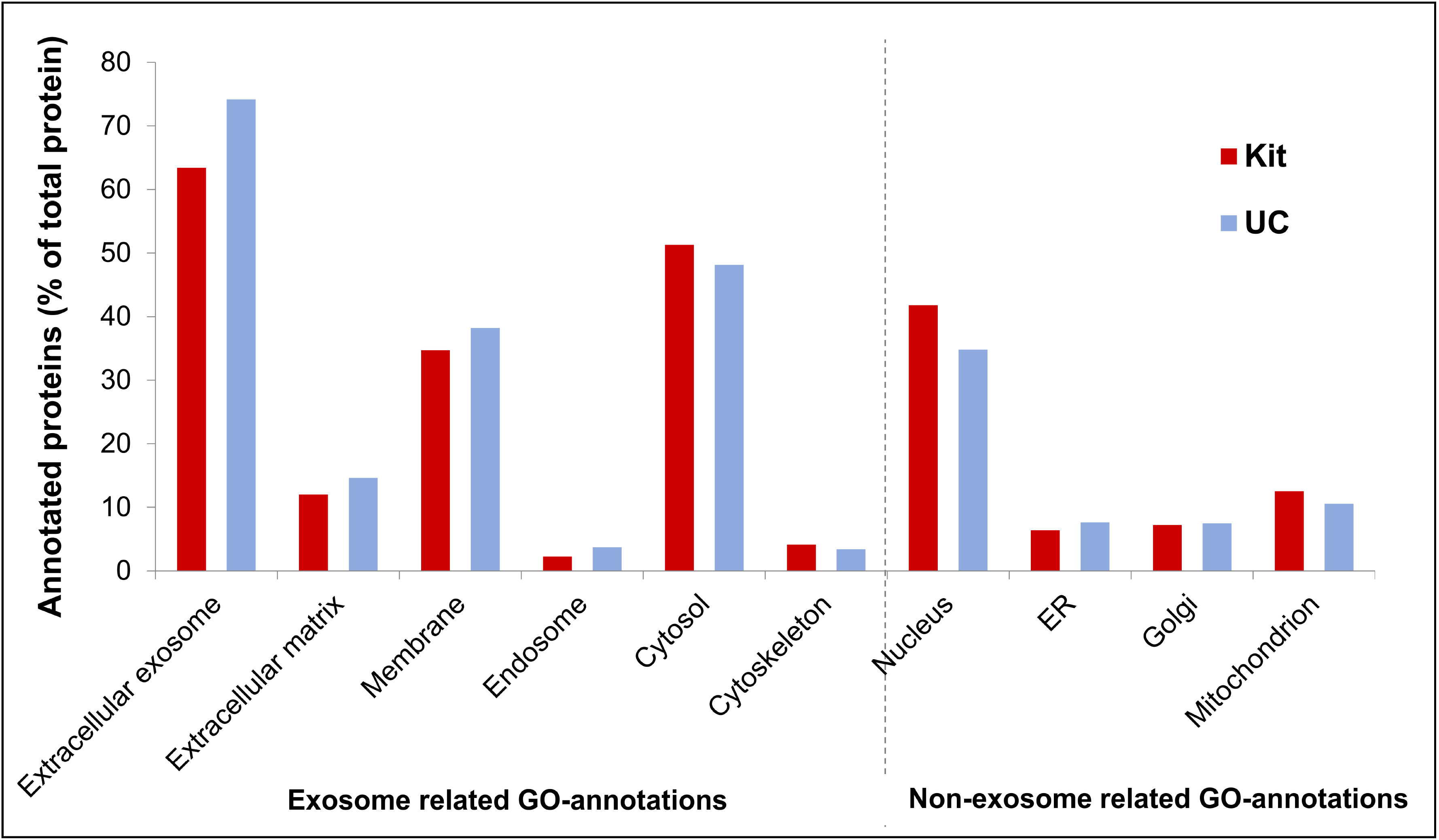
Chromatograms and MS/MS spectrums from LC-MS/MS analysis of GBM- and BC exosome peptides. **A**) Chromatogram with corresponding MS/MS spectrum for the CD9 signature peptide KDVLETFTVK (*m/z*=393.89, *z*=3) in BC exosomes isolated by UC. C) Chromatogram with corresponding MS/MS spectrum for the calnexin signature peptide AEEDEILNR (*m/z*=544.77, z=2) from GBM exosomes isolated by UC. An in-house packed 50 μm x 150 mm column with 80 Å Accucore particles with C_18_ stationary phase was used for separation. A 50 μm x ~3 mm in-house packed pre-column with the same column material was used for trapping. The elution was performed with a linear gradient of 3-15 % MP B in 120 minutes. See **Section 2.11.1** for more LC-MS/MS parameters.

LC-MS/MS could also identify a number of positive markers (see **Figure 5** for examples). However, there was expectedly not a complete overlap with those observed with WB, as e.g. sensitivity can vary between WB and untargeted LC-MS/MS. In-house prepared nanoLC columns packed with core shell particles provided high-resolution separations (**Figure 5**, and see reference [57] for packing procedure). Examples of potential biomarkers for GBM, e.g. heat shock proteins 70 kDa and 90 kDa [58–60], chondroitin sulfate proteoglycan 4 [58,61], CD44 [62,58,61] and CD276 [63] were identified using LC-MS/MS. Examples of LC-MS/MS-detected biomarkers related to triple negative breast cancer were e.g. heat shock 90 kDa *α* and β protein [64], calmodulin and epithermal growth factor receptor [65] (see **Supplemental Proteins**). When comparing cell sources, the number of identified proteins was lower in GBM isolates than BC isolates, but the number of identified proteins for GBM isolates was comparable to another LC-MS/MS study on GBM exosomes [66].

**Figure 5.**
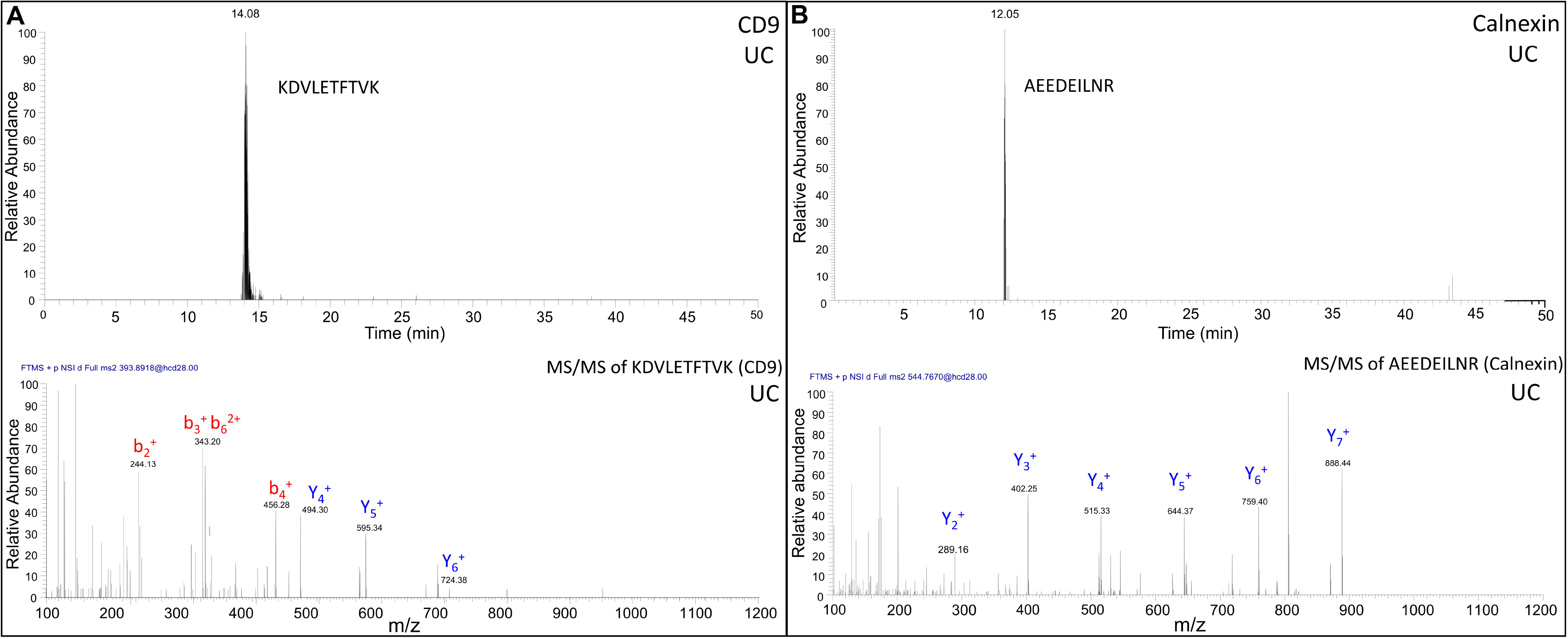
GO annotation of proteins in BC exosomes to different cellular locations. The identified proteins classified by their cellular location (GO annotations) grouped based on their positive/ negative relevance towards exosomes. The annotated proteins (% of total proteins) and their cellular location, with proteins annotated from the kit isolates are shown in red (from 749 DAVID ID’s), while proteins annotated from the UC isolates are shown in blue (from 615 DAVID ID’s).

## 4 Conclusions

Regarding our glioblastoma/breast cancer “case study” samples, the UC/kit isolation methods overall were approximately equal in quality. Kit isolation however has an advantage of requiring less starting material compared to conventional UC equipment. Untargeted LC-MS/MS revealed a number of biomarkers related to the diseases, supporting the concept of exosomes being an interesting matrix towards diagnostics. In addition to exosomes, our analyses suggest the presence of cellular contaminations and other vesicles. Hence, the “isolations” should perhaps be considered “enrichments”. Considering that the methods do not fully provide isolations, we welcome alternative approaches to preparing and analyzing these important extracellular vesicles.

## Acknowledgments

This work was supported by the Department of Chemistry, University of Oslo, Norway. We would like to acknowledge DIATECH@UiO, since parts of this work have been carried out within this strategic research initiative at the Faculty of Mathematics and Natural Sciences, University of Oslo. This work has also been supported by the UiO:Life Science funded convergence environment “Organ on a chip and nano-devices”. Conflict of Interest: The authors declare that they have no conflict of interest.

---

**Figure 6 Venn diagram presenting the number of proteins identified by LC-MS/MS in exosomes isolated by kit and UC from GBM- and BC cell culture medium.** The numbers are the total number of unique proteins identified when trypsin, keratin related proteins and the proteins identified in blank isolates were disregarded. One signature peptide was selected as requirement for positive identifications during database search. Equal amounts of protein were injected for both kit- and UC isolates (~ 1.5 μg protein for GBM isolates (n = 6) and ~2-5 μg protein for BC exosomes (n=3)). A list of all proteins identified is presented in **Supplemental Proteins**.

